## Supplementary material for "A two-step mechanism for creating stable, condensed chromatin by the Polycomb complex PRC1": sup_figures

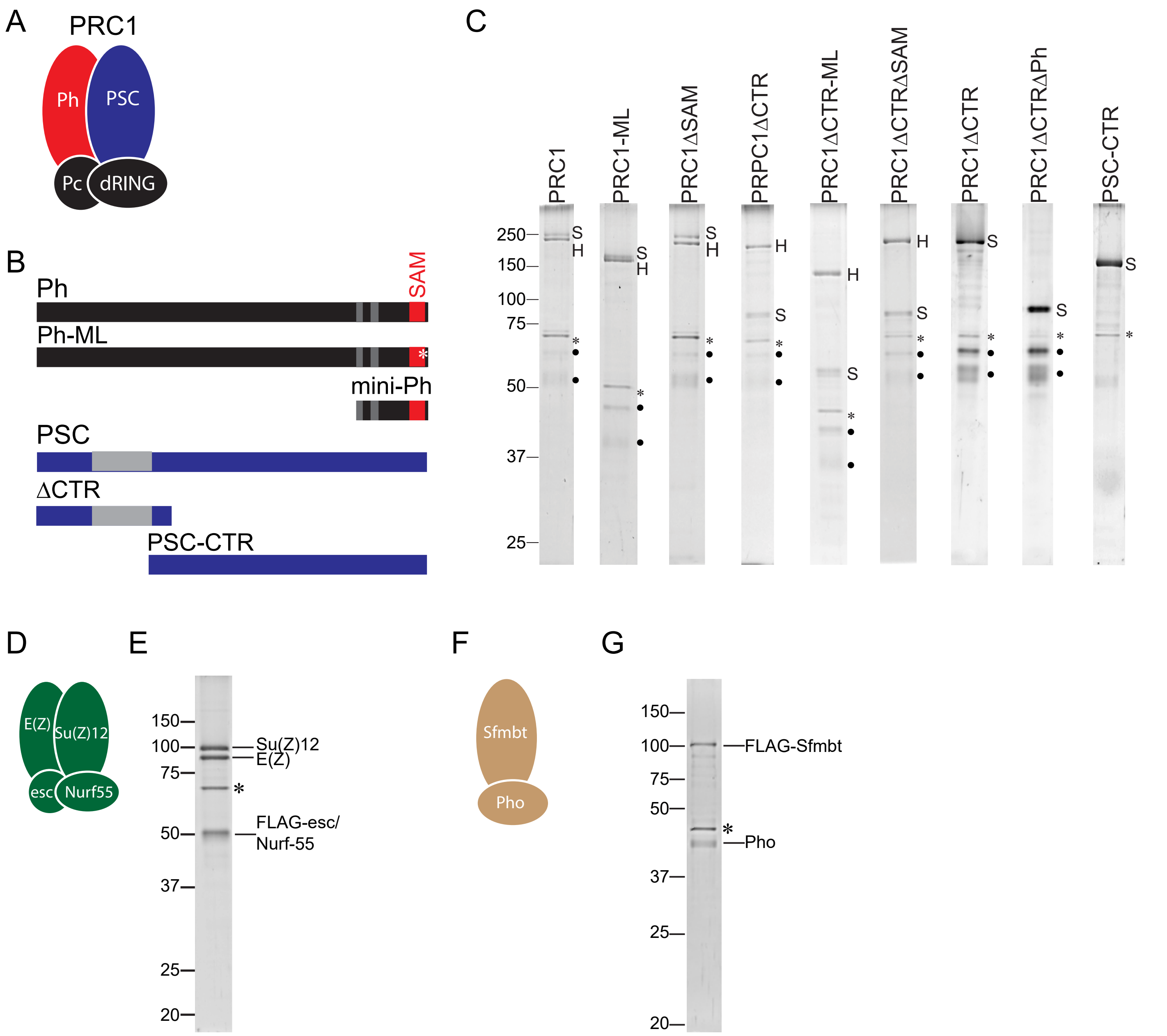

A
PRC1
Ph
PSC
Pc
dRING
B
Ph
SAM
Ph-ML
mini-Ph
PSC
ΔCTR
PSC-CTR
C
PRC1
PRC1-ML
PRC1ΔSAM
PRPC1ΔCTR
PRC1ΔCTR-ML
PRC1ΔCTRΔSAM
PRC1ΔCTR
PRC1ΔCTRΔPh
PSC-CTR
250
150
100
75
50
37
25
S
H
D
E(Z)
Su(Z)12
esc
Nurf55
E
150
100
75
50
37
25
20
Su(Z)12
E(Z)
FLAG-esc/
Nurf-55
F
Sfmbt
Pho
G
150
100
75
50
37
25
20
FLAG-Sfmbt
Pho

**Figure S1 Protein preparations.** A. Schematic of PRC1. B. Domain organization of Ph and PSC and truncations/mutations used. Gray regions and the SAM are structured domains; the rest of both proteins is predicted or shown to be disordered. C. SDS-PAGE gel stained with SYPRO Ruby showing different PRC1 variants of Ph and/or lacking the PSC-CTR, and the PSC-CTR alone. S indicates the position of PSC, H the position of Ph. Asterisk is co-purifying Hsc70 and filled circles indicate Pc (top) and dRING (bottom). Lanes are from different gels; marker is only shown for the first lane. D, E. Schematic (D) and SYPRO Ruby-stained SDS-PAGE (E) of PRC2. F, G. Schematic(F) and SYPRO Ruby stained SDS-PAGE (G) of PhoRC. Asterisk in E and F indicates co-purifying Hsc70.

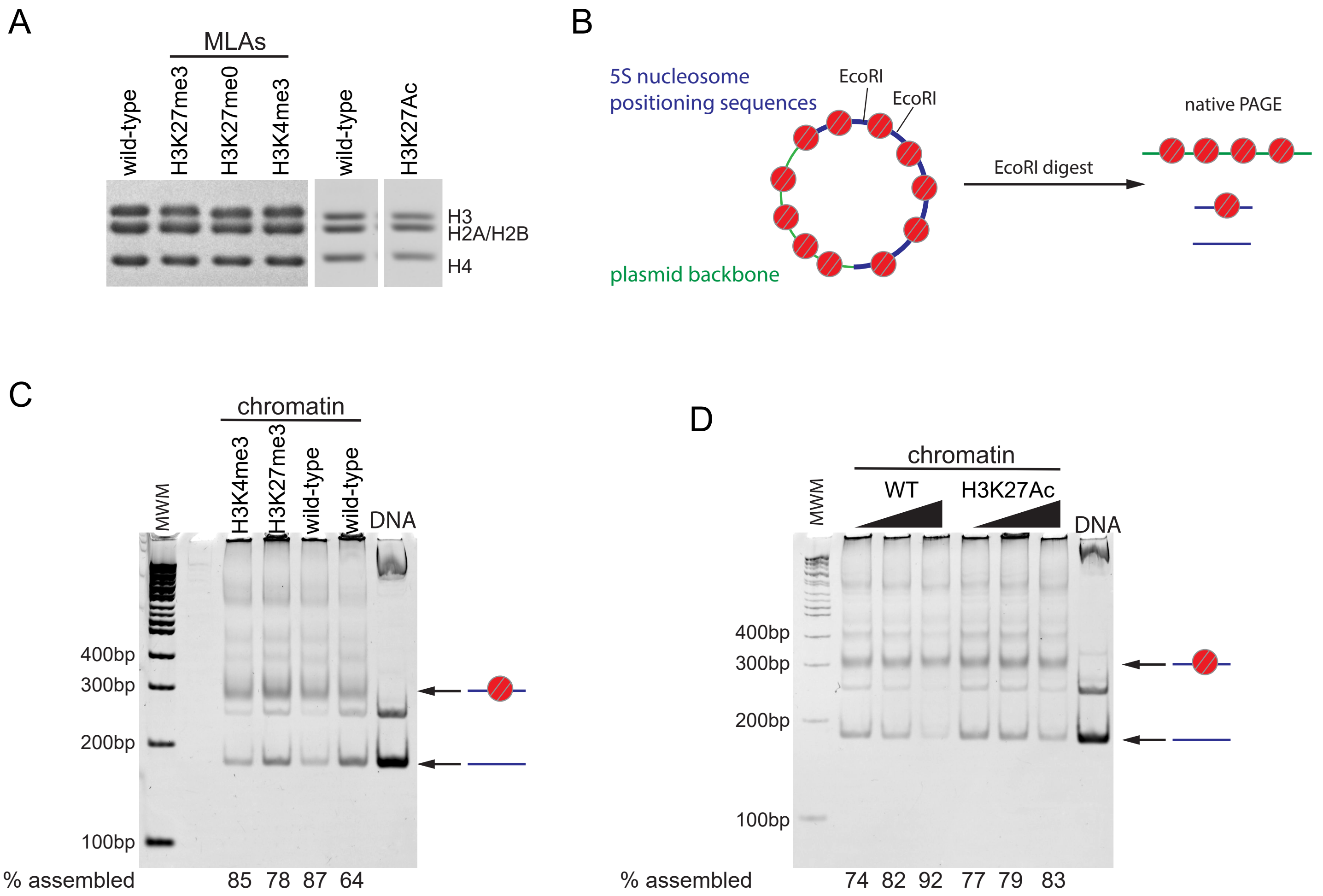
A
MLAs
wild-type
H3K27me3
H3K27me0
H3K4me3
wild-type
H3K27Ac
H3
H2A/H2B
H4
B
5S nucleosome
positioning sequences
EcoRI
EcoRI
plasmid backbone
EcoRI digest
native PAGE
C
chromatin
MWM
H3K4me3
H3K27me3
wild-type
wild-type
DNA
400bp
300bp
200bp
100bp
% assembled
85 78 87 64
D
chromatin
WT
H3K27Ac
MWM
DNA
400bp
300bp
200bp
100bp
% assembled
74 82 92 77 79 83

**Figure S2 Chromatin preparation.** A. Coomassie blue stained SDS-PAGE of reconstituted histone octamers with the indicated modifications. B. Schematic of the EcoRI digest assay to measure nucleosome assembly over the 5S repeats (taken from [35]). C, D. EcoRI analysis of wild type and MLA-containing chromatin (C), and wild type and H3K27Ac chromatin (D).

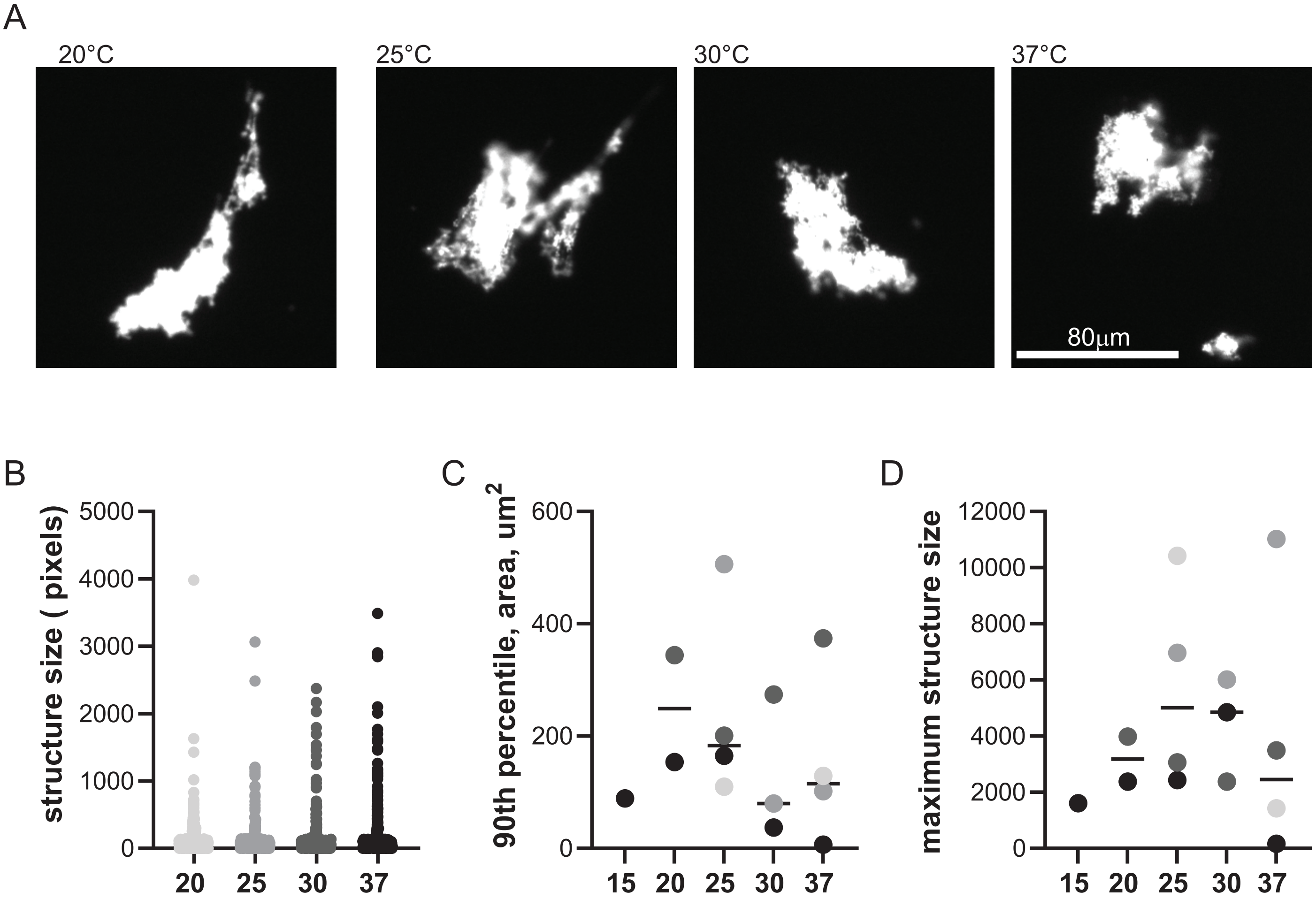
A
20°C
25°C
30°C
37°C
80μm
B
structure size ( pixels)
5000
4000
3000
2000
1000
0
20
25
30
37
C
90th percentile, area, um2
600
400
200
0
15
20
25
30
37
D
maximum structure size
12000
10000
8000
6000
4000
2000
0
15
20
25
30
37

**Figure S3 Effect of temperature on PRC1-chromatin condensate formation**. A. Images of structures formed by PRC1 + chromatin during overnight incubation at indicated temperatures. B. Quantification of structures formed at different temperatures for a representative experiment. No differences between pairs of samples (i.e. 20-25, 25-30, 30-37) were detected by Kruskal-Wallace test with Dunn's correction for multiple comparisons. C, D. Summary of structures formed at different temperatures, showing the 90$^{th}$ percentile (C) and maximum (D) area of structures formed. Symbols that are the same shade are from the same experiment.

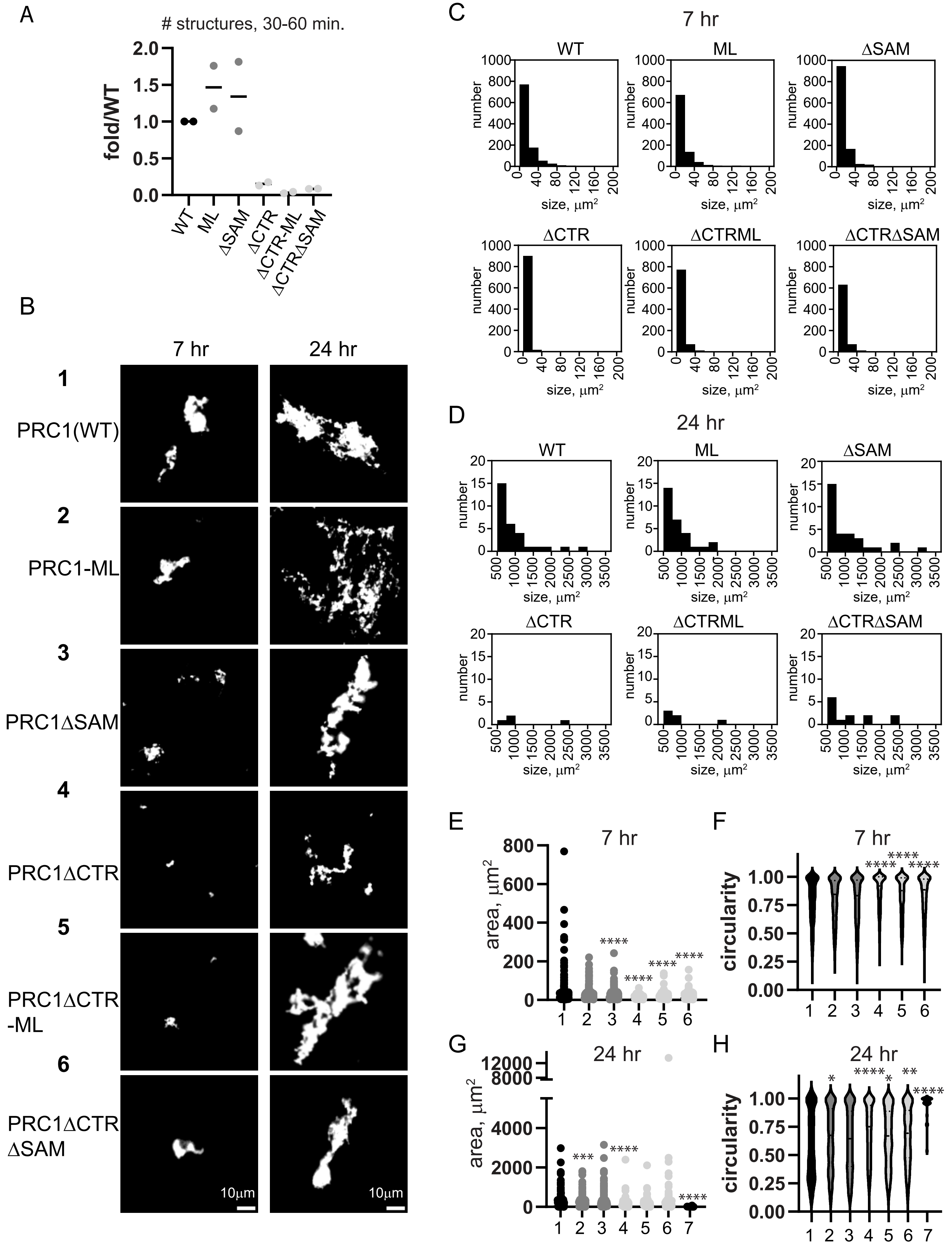
A
### structures, 30-60 min.
fold/WT
2.0
1.5
1.0
0.5
0.0
WT
ML
ΔSAM
ΔCTR
ΔCTR-ML
ΔCTRΔSAM
B
7 hr
24 hr
1
PRC1(WT)
2
PRC1-ML
3
PRC1ΔSAM
4
PRC1ΔCTR
5
PRC1ΔCTR
-ML
6
PRC1ΔCTR
ΔSAM
10μm
10μm
C
7 hr
WT
ML
ΔSAM
ΔCTR
ΔCTRML
ΔCTRΔSAM
number
size, μm²
0
40
80
120
160
200
1000
800
600
400
200
D
24 hr
WT
ML
ΔSAM
ΔCTR
ΔCTRML
ΔCTRΔSAM
number
size, μm²
500
1000
1500
2000
2500
3000
3500
20
15
10
5
0
E
7 hr
area, μm²
800
600
400
200
0
1
2
3
4
5
6
****
****
****
****
F
7 hr
circularity
1.00
0.75
0.50
0.25
0.00
1
2
3
4
5
6
****
****
****
G
24 hr
area, μm²
12000
8000
4000
2000
0
1
2
3
4
5
6
7
***
****
****
H
24 hr
circularity
1.00
0.75
0.50
0.25
0.00
1
2
3
4
5
6
7
*
****
*
**
****

**Figure S4 The PSC-CTR increases the rate of formation and size of PRC1-chromatin condensates.** A. Quantification of structures formed at short time points (30 or 60 min.) by different complexes in two different experiments. Numbers were normalized to the number in reactions with wild-type PRC1. B. Representative structures formed with different complexes after 7 or 24 hours. C. Histograms of the number structures formed by different complexes after 7 hours of incubation. D. Histograms showing the number of structures greater than 500$\mu m^2$ formed by different complexes with chromatin after 24 hours. Chromatin graph is not shown as no structures in this size range were observed. E-H. Quantification of area (E, G) or circularity (F, H) of structures formed by different complexes at different time points. Asterisks are for Kruskal-Wallace test with Dunn's correction for multiple comparisons (****=$p \leq 0.0001$). See **Table S1** for a summary of comparisons of area and circularity across multiple experiments.

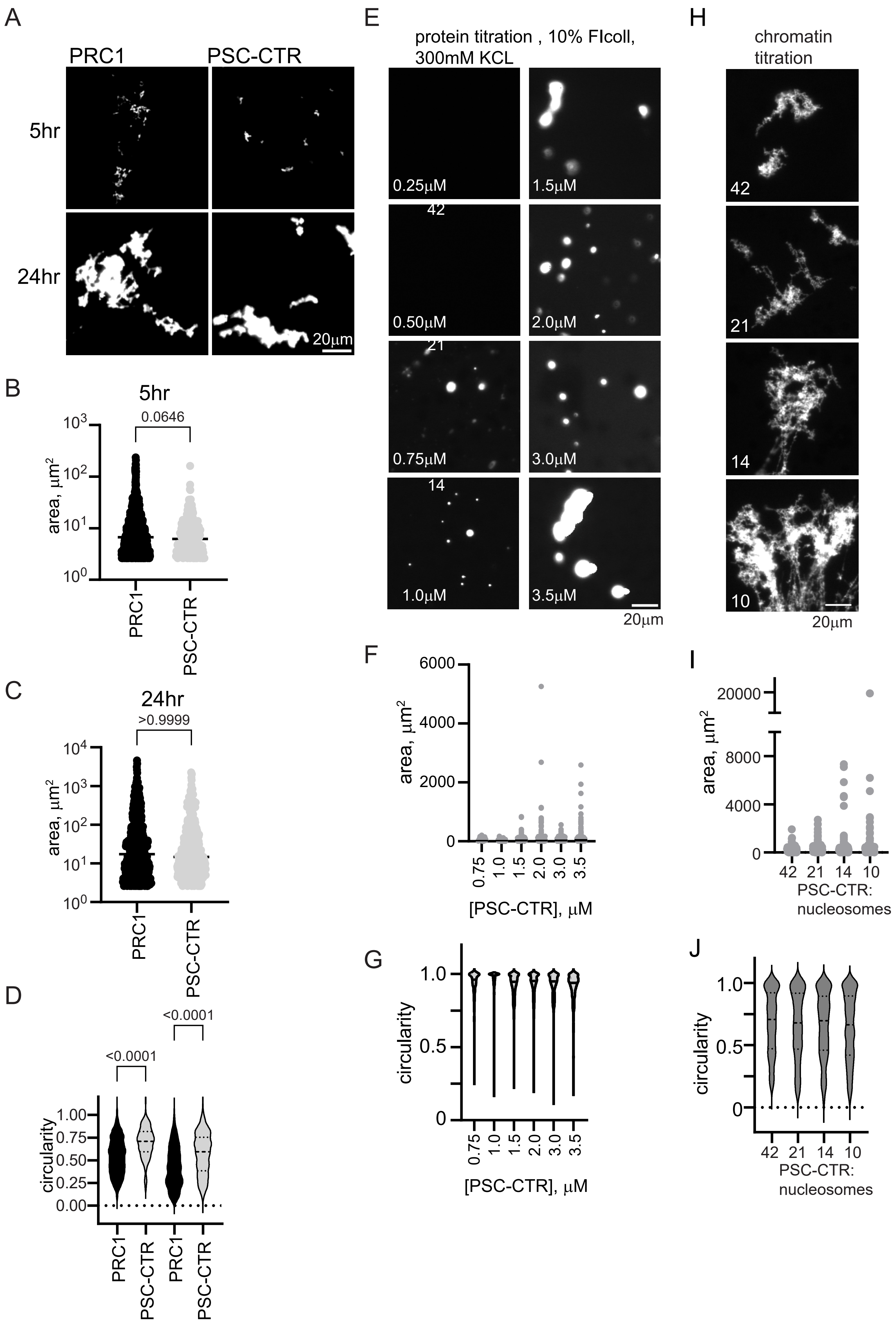
A
PRC1
PSC-CTR
5hr
24hr
20µm
B
5hr
0.0646
area, µm²
PRC1
PSC-CTR
C
24hr
>0.9999
area, µm²
PRC1
PSC-CTR
D
<0.0001
<0.0001
circularity
PRC1
PSC-CTR
PRC1
PSC-CTR
E
protein titration , 10% FIcoll, 300mM KCL
0.25µM
1.5µM
0.50µM
2.0µM
0.75µM
3.0µM
1.0µM
3.5µM
20µm
F
area, µm²
[PSC-CTR], µM
G
circularity
[PSC-CTR], µM
H
chromatin titration
42
21
14
10
20µm
I
area, µm²
PSC-CTR: nucleosomes
J
circularity
PSC-CTR: nucleosomes

**Figure S5 The PSC-CTR forms round condensates with Ficoll, and large networks with chromatin at 120mM KCl.** A. Representative images of structures formed by PRC1 (10nM) or the PSC-CTR (80nM) with chromatin (replicate of experiment shown in **Figure 4A-D**). B-D. Quantification of structure areas after 5 (B) or 22 (C) hours, and of circularity (D) for both time points. E. Condensates formed by the PSC-CTR in the presence of Ficoll (100mg/ml). F, G. Quantification of area (F) and circularity (G) of titration shown in E. H. Titration of chromatin with the PSC-CTR (0.72µM). Number indicates the ratio of protein to nucleosomes, which were used at ~17, 34, 52, and 69nM. I, J. Quantification of area (I) and circularity (J) of titration shown in H, indicating that structures increase in size with increased addition of chromatin.

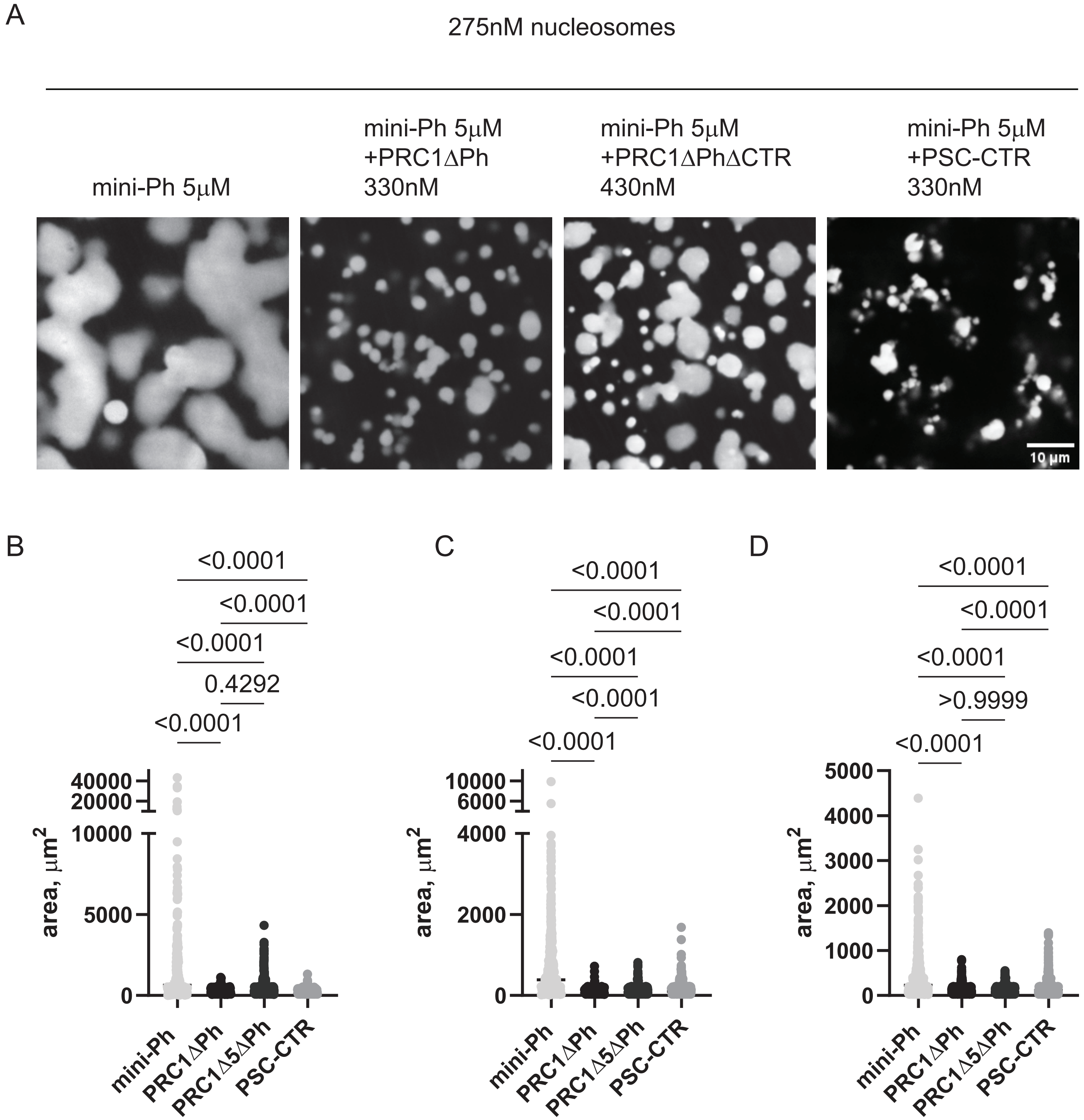
A
275nM nucleosomes
mini-Ph 5μM
mini-Ph 5μM +PRC1ΔPh 330nM
mini-Ph 5μM +PRC1ΔPhΔCTR 430nM
mini-Ph 5μM +PSC-CTR 330nM
10 μm
B
C
D
<0.0001
0.4292
>0.9999
area, μm2
mini-Ph
PRC1ΔPh
PRC1Δ5ΔPh
PSC-CTR

**Figure S6 PRC1 lacking the PSC-CTR requires higher concentrations to arrest mini-Ph-chromatin condensates.** A. Representative images of reactions with 5mM mini-Ph and 275nM nucleosomes after addition of buffer, PRC1ΔPh, PRC1ΔCTRΔPH, or the PSC-CTR at the indicated concentrations. B-D. Quantification of the area of condensates in three replicates of the experiment shown in A. p-values are for Kruskal-Wallace test with Dunn's correction for multiple comparisons.

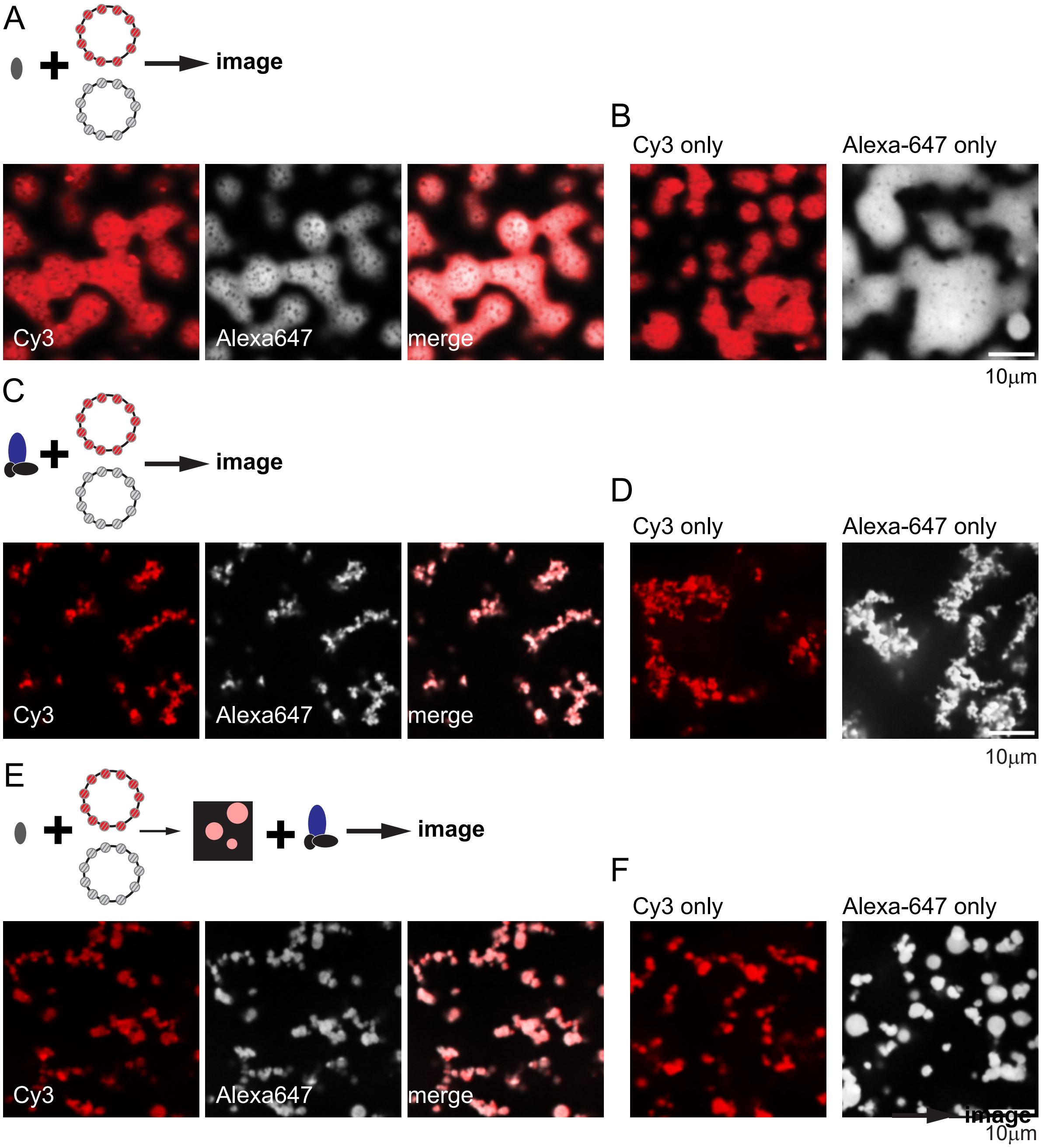
A
image
B
Cy3 only
Alexa-647 only
Cy3
Alexa647
merge
10μm
C
image
D
Cy3 only
Alexa-647 only
Cy3
Alexa647
merge
10μm
E
image
F
Cy3 only
Alexa-647 only
Cy3
Alexa647
merge
image
10μm

**Figure S7 Control experiments for mixing experiments with different chromatin templates.** A, C, E show representative images of reactions where both templates are added prior to adding the indicated protein. B, D, F show images of reactions with single template.

A

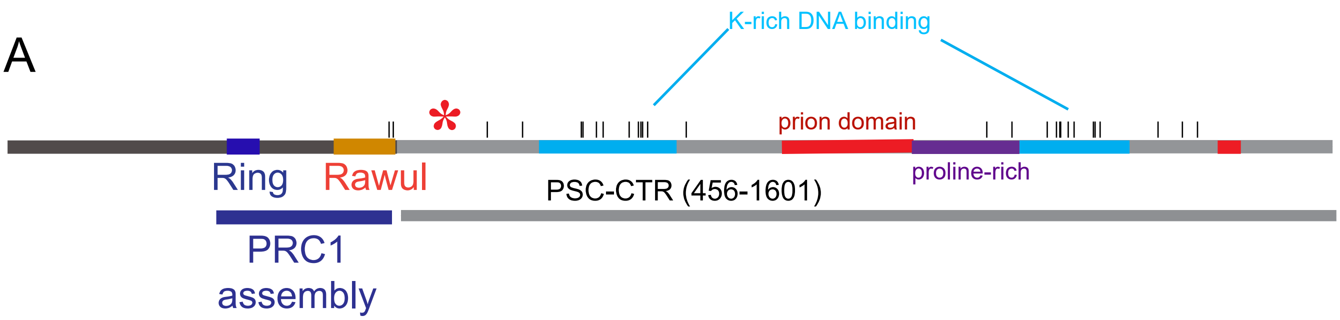


B sequence of prion domain:

small non-polar
small polar
aromatic
positively charged
negatively charged
proline

PNSPIYSPSS PQYVPSYNIP TMPTYKYTPK PTPNSGSGNG GSGSYLQNML
GGGNGGSLGG LFPSPPTKSD QNTNPAQGGG GSSSATQSGG NNGIVNNNIY
MPN

**Figure S8 Additional analysis of the PSC-CTR.** A. Sequence and functional features in the PSC-CTR. The two putative DNA binding regions were identified in protein footprinting experiments [46], while proline rich and prion domains were identified by sequence analysis. PLAAC [57] was used to identify the prion domains. The region between the two DNA binding domains forms many contacts with other parts of PSC in cross-linking mass spec experiments [46]. Black lines indicate phosphorylation sites identified in our AP-MS analysis (although not independently validated) or available on UniProt. Asterisk indicates position of a truncation mutation that is lethal in *Drosophila* embryos [18, 42]. B. Sequence of the main prion domain.
